# Ecological stoichiometric characteristics of Carbon (C), Nitrogen (N) and Phosphorus (P) in leaf, root, stem, and soil in four wetland plants communities in Shengjin Lake, China

**DOI:** 10.1101/2020.02.24.962456

**Authors:** Dagne Tafa Dibar, Kun Zhang, Suqiang Yuan, Jinyu Zhang, Zhongze Zhou, Xiaoxin Ye

**Author notes:** Corresponding author: Dagne Tafa, Email address.

## Abstract

Ecological stoichiometric should be incorporated into management and nutrient impacted ecosystems dynamic to understand the status of ecosystems and ecological interaction. The present study focused on ecological stoichiometric characteristics of different macrophyte plants soil, leave, stem and root after the removal of seine fishing since 2000 from Shengjin Lake. For C, N and P analysis from leaves, stems, roots and soil to explore their stoichiometric ratio and deriving environmental forces here four dominant plant communities (*Zizania caduciflora, Vallisineria natans, Trapa quadrispinosa* and *Carex schmidtii*) were collected. C, N, P and C: N: P ratio in leafs, stems, roots and soil among the plant communities vary and the studied plant communities had significant effect on the measured variables. There was high C: N in *C.schmidtii* soil (7.08±1.504) but not vary significantly (*P* >0.05), and N: P ratio measured high in *V. natans* (13.7±4.05) and C: P in *T.quadrispinosa soil* (81.14±43.88) and showed significant variation (*P*<0.05) respectively. High leaf C: N and N: P ratio was measured in *C. schmidtii* and *V. natans* respectively. Nevertheless, high leaf C: P ratio was measured in *Z.caduciflora*. From the three studied organs leafs C: N, N: P ratio showed high values compared to root and stems. The correlation analysis result showed that, at 0-10cm depth ranges SOC correlated negatively with stem total phosphorus (STP), and RTN (*P*<0.05) but positively strongly with LTP and LTN (*P*<0.01) respectively. Soil total nitrogen at 0-10cm strongly positively correlated with LTP (*P*<0.01) and positively with RN: P and LTC (*P*<0.05). Soil basic properties such as SMC.BD and pH positively correlated with soil ecological stoichiometric characteristics. Redundancy analysis (RDA) result showed pH and available phosphorus were the potential determinant of soil stoichiometry.

## Introduction

Ecological stoichiometry is an important tool for studying ecological processes and functions (Shang B *et al*., 2018). Besides, it uses to explore the dynamic balance of various elements and their interactions (Moorthi S D *et al*., 2016) and limitation of plant growth (Chen *et al*., 2016). Carbon (C), Nitrogen (N) and phosphorus (P) concentrations are vital element sources in plant and their changes in characteristics limit plant growth (Sperfeld E.et *al*., 2017). Further, Nitrogen (N) and Phosphorus (P) elements are determinant nutrients for plant growth and aquatic ecosystems functioning (Elser *et al*., 2007) and their mass ratio can potentially affect plant-mediated ecological processes (Zechmeister *et al*., 2015). Soil C: N: P ratios can directly reflect soil fertility and indirectly the nutritional status of plants and species composition of plant communities (Mao *et al*., 2016). Carbon, nitrogen, and phosphorus biogeochemical circulations closely related to the soil’s ecological structure, processes, and functions in wetland ecosystems (Hansson *et al*., 2005). However, it varies due to the difference in vegetation characteristics (Sardans and Penuelas, 2014), plant identity associated with growth rate, and nutrients allocation (Yu *et al*., 2012), and plants size, taxa and life forms (Zhang.Z. *et al*., 2013). Recently stoichiometry of C: N: P has been applied to understand nutrients limitation, community dynamism (Johnson and Agrawal, 2005), nutrient use efficiency (He *et al*., 2010) and the global biogeochemical cycle (Schmidt *et al*., 2016) in both terrestrial and aquatic ecosystems. Moreover, plant C: N: P stoichiometry is strongly influenced by nutrient availability, and can effectively show the changes in C, N, and P cycles (Hessen *et* al.,2013), control plant functional type, climate and anthropogenic interference (Long *et al*., 2016). Ecological stoichiometry especially plants leaf C, N, and P play a vital role in analyzing the composition, structures, and function of the concerned community and ecological systems (Gao *et al*., 2013; Zhang *et al*., 2013). Under different environmental conditions, plant physiological processes (Liu *et al*., 2015), wetland hydrology, soil pH (Sato and comerford, 2005) and salinity (Sun *et al*. 2013), and community type (Shang *et al*., 2013) can determine these nutrients and in turn it can be used for the accumulation and allocation of plant biomass (Lie and Li, 2016). Because of human interference (Sardans and Penuelas, 2014), climate change, hydrological fluctuation, aquatic and offshore environment wetlands have been greatly affected. This is posing great impacts on vegetation and ecosystems (Qin et al.2014). C: N and N: P ratio can be used as effective indicators for the health conditions and growth status of plants (Vrede et al., 2004; Hessen et al., 2007) and understating the life strategies of plants (Wu et al., 2010). Therefore, analysis of C, N and P ecological stoichiometry and their corresponding ratio C: N: P are important. This mainly provides good information to know different plants adaptation capacity to changing environment and stress. Besides, it helps to design, formulate strategies and policy for environmental protection and ecological restoration programs. Submerged plant communities have dominated our study site before 2000 (Meng, 1979). However, due to over fish production and using seine-fishing net they have been drastically affected. Thus, the government officially banned seine fishing since 2008 and those drastically disappeared vegetation’s began restoring gradually. To our knowledge, there was no study conducted on the ecological stoichiometry of wetland soil in this Lake after the regeneration of those degraded plants. With this knowledge gap, this study mainly sought to determine, (1) the distribution patterns and stoichiometric characteristics of C,N, P and C:N:P in soil, leave, stem and root of different macrophytes communities (2) to analyse soil-plant ecological stoichiometric interaction.

## Materials and Methods

### Study site description

Shengjin Lake is located at (30°16’–30°25’N, 116°59’-117°12’E), in the southern bank of the Yangtze River, close to Anqing (Fig. 1). It has an area of 133 km^2^ in the wet and 33 km^2^ in the dry seasons respectively. The protected area is centered on Shengjin Lake and the coast extends outward by about 2.5 km^2^. The maximum area of the lake during the flood peak is ~14000 ha (17.0 m (asl) but, the water level normally falls each year to less than 10 m (asl) during the dry seasons and decrease to ~3400 ha. It gets inflow from three small rivers flowing directly into the lake and from the Yangtze River via the Huangpen Sluice. Various terrains with low mountains, hills in the southeast and plains in the northwest surround the lake.

**Fig.1.**
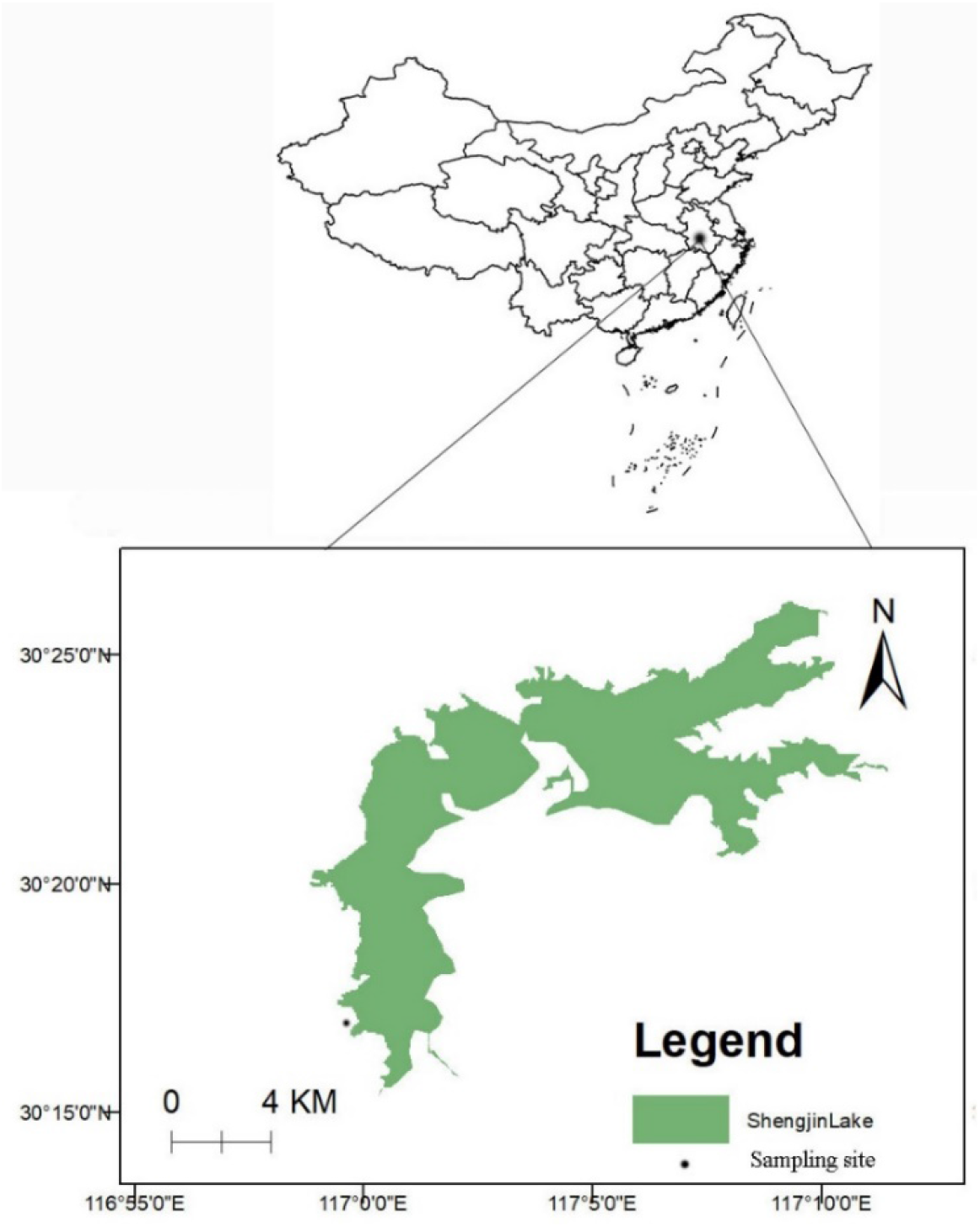
Sampling site geographical location

### Soil and plant sampling

In order to examine ecological stoichiometry of C, N and P in soil, the soil samples were collected from four macrophyte plants i.e *Zizania caduciflora, Vallisineria natans, Trapa quadrispinosa* and *Carex schmidtii* respectively. To fix the area of sampling plot 1m×1m framed quadrat was used and the sample were collected using a 5cm diameter soil auger in October 2019 randomly. Soil sampling profile sectioned in to 10cm interval as 0-10cm, 10-20cm and 20-30cm in depth vertically after (Wu *et al*., 2012a). In each depth and sampling plots, three replicate samples were taken and mixed evenly to homogenize and composited. The homogenized soil samples were grouped per specific plant communities, placed in polyethylene plastic bag, and transported to the laboratory for analysis. For plant C (PTC), (PTN) and (PTP) analysis, we collected root, stem and leave. The root samples were uprooted properly and cut off with scissors, washed with hot tap and distilled water for analysis in the laboratory (Pregitzer *et al*., 2002). The leaves and stems surface were washed by tap and distilled water in the laboratory to avoid epiphytic and adhering muds to the surface.

### Laboratory analysis

At room temperature, both soil, plant leave, stem and root samples were air dried in open space. From the homogenized soil samples visible stones, rocks, shell, plant debris, and roots were removed by hand. The air dried soil samples were crushed and ground using mortar and pistil and sieved through 0.15mm sieve before analysis. Leave samples (without petiole and rachis), stem and root were ground using a ball mill after (Cong *et al*., 2014) and sieved through 0.15mm sieve to analyse carbon, total nitrogen, and total phosphorus. All samples were triplicated and averaged procedurally during analysis. Carbon contents in soil, leaf, stem and root were treated by wet oxidation of organic matter with potassium dichromate (K_2_Cr_2_O_7_) solution and sulfuric acid (H_2_SO_4_), followed by ferrous sulfate (FeSo_4_) as titrant (Zhang *et al*., 2015). Analysis of TN from soil, leaf, stem, root, and available nitrogen (AN) was carried out using perchloric acid (HClO_4_) by digesting with sulfuric acid (H_2_SO_4_) and measured by UV-2450 spectrophotometer (Shimadzu Scientific Instruments, Japan) (Romanya *et al*., 2017). After digesting by sulfuric acid (H_2_SO_4_) and hydrogen oxide (H_2_O_2_) a TU-1901DS ultraviolet spectrophotometer UV-2300 (Tec comp Com, Shanghai, China) molybdenum antimony colorimetric method was used to quantify soil, leaf, stem and root total phosphorus (TP) and available phosphorus (AP)) (Zhang *et al*., 2015). Finally, C, N and P concentrations were presented as g kg-^1^ on a dry mass basis (Lu *et al*., 2012) and available (AN) and available phosphorus(AP) in (mg kg-^1^). The stoichiometry of C: N: P molar ratios was computed as mass ratio proportion. Portable pH meter (Sensor, Hach, USA) was used to measure soil pH and Electrical conductivity (EC) after the samples have been made 1:5 soil/water ratio in deionized water in the suspension after mixing the samples for one hour intermittently (Estefan et al.2013).Barua and Barthakur, 1999). Soil moisture contents (SMC) and bulk density (BD) were determined by the oven-dry method (Zhang *et al*., 2015).

### Quality assurance

To ascertain our experimental quality at the time of sample digestion and processing, we used the blank sample to manage the background effect of materials used for each analyzed samples. Besides, all used sampling bottles have been soaked in acid solution (HCl) from 30-40 minutes, washed in deionized water and oven dried before use accordingly.

### Data analysis

One way analysis of variance (ANOVA) was applied for the statistical significance test. Mean and the standard deviation was used to describe all variables in the statistical analysis and mean values were reported at 95% confidence interval. Person correlation coefficient analysis was used to test the relationship between TOC, TN, TP, and C: N: P ratios in leave, stem, root, soil and environmental variables in different plant communities at (p<0.05). The data were tested for normality distribution and homogeneity of variance before parameter test. SPSS.20 (IBM Corporation, Armonk, New York) software was used for all statistical analysis. RDAwas carried out using Canoco 4.5 for Windows (Microcomputer Power, Ithaca, NY) to select the best explanatory variables. The data and Monte Carlo reduced model tests with 499 unrestricted permutations were used to statistically evaluate significance. All columnar figures were done using Graph Pad Prism 5 (https://www:graphpad.com)software) software.

## Results

### Soil C, N and P distribution patterns among different plant communities

Different vegetation types had significantly affected soil organic carbon (SOC), soil total (STN) and soil total phosphorus (STP) distribution and varied significantly (*P* <0.05) (Fig.2A, B, C). As illustrated in (Fig.2A), SOC in plants communities varied from 32.6±4.59g kg^-1^_12.56±0.342 g kg^-1^. Among the plant communities this is ordered as *Carex schmidtii* > *Zezinia caduciflora* > *Vallisineria natans* > *Trapa quadrispinosa* respectively. The range of STN found higher in *C. schmidtii* (10.18±2.56 g kg^-1^) followed by *V. natans* and *Z. caduciflora* (6.953±1.23g kg^-1^) and (4.0.74±0.734 g kg^-1^) from 0_10cm soil layer respectively (Fig.2B). The STP values along with the depth profiles were ranged from 0.814±0.3302 g kg^-1^ - 0.1594±0.1581 g kg^-1^ (Fig.1C). Among the four plant communities, high SOC values (32.6±4.59 g kg^-1^) and STN (10.18±2.56 g kg^-1^) were measured in *C. schmidtii* and showed significant difference (*P* <0.05), (Fig.2A, B). High STP values were measured in *V. natans* especially at 0-10cm depth ranges and differed significantly (*P* <0.05), (Fig. 2C).

**Fig. 2.**
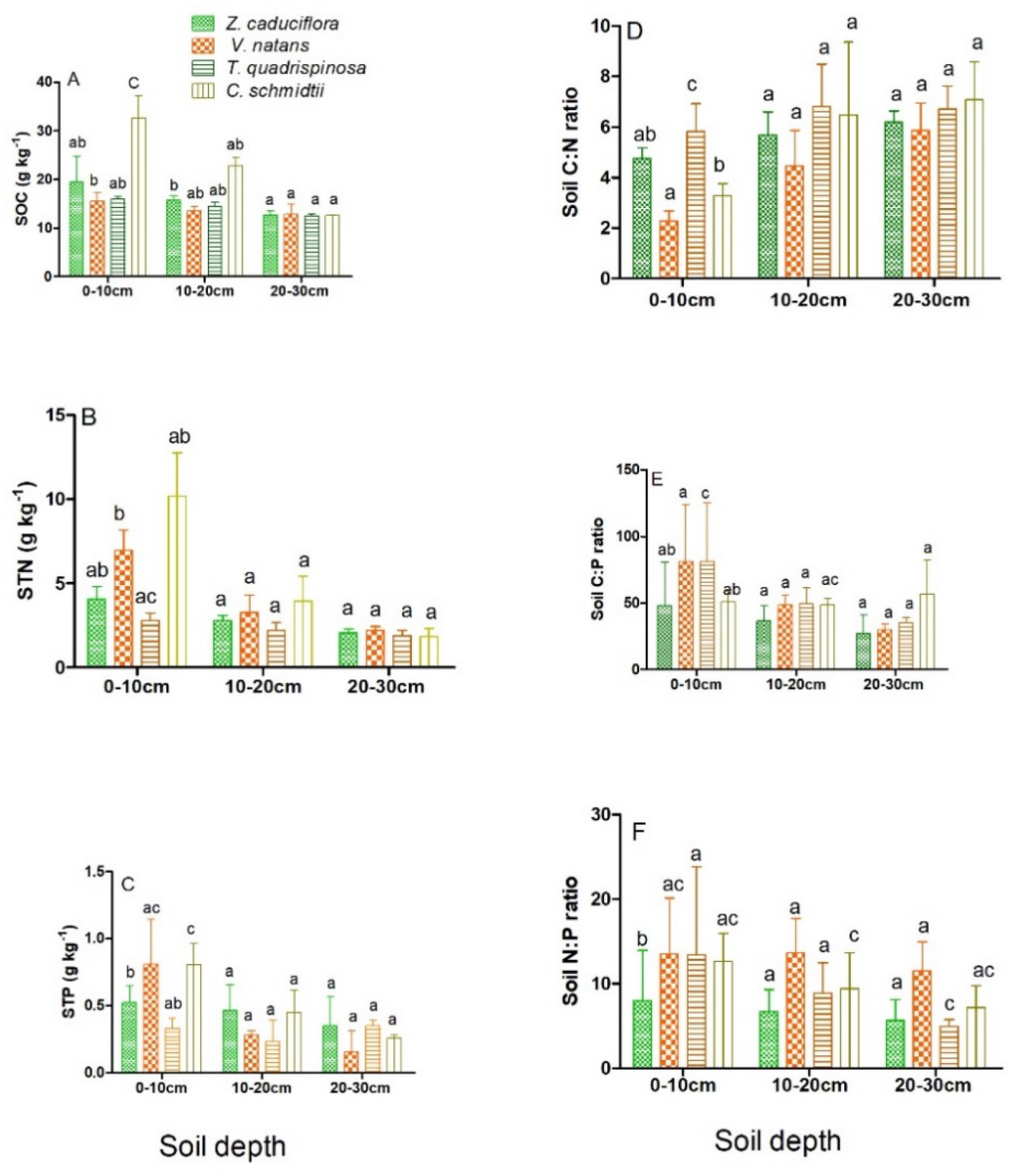
The vertical distribution of SOC(A),STN(B),STP(C),C:N(D),C:P(E), N: P(F). Note: The values were displayed by mean±sdav in each depth profiles. Mean denoted by different letters shows significant difference (P <0.05), based on ANOVA, LSD and Tukey result (2-tailed) analysis. SOC, soil organic carbon, STN, soil total nitrogen, STP, soil total phosphorus.

### Soil C: N, C: P, N: P ratios distribution pattern in four plant communities vertically

Ecological stoichiometry of C: N, C: P and N: P ratios in different plant communities were found vary significantly. The distribution of C: N ratio was varied from 2.28±0.392 - 7.084±1.504 (Fig.2D). C: N ratio displayed slightly high in *C. schmidtii* and showed peak value (7.08±1.504), but showed no significant difference (*P* >0.05), (Fig.2D). In the same mode C: P and N: P ratios were ranged from 27.48±13.78 −81.5±43.88, while N: P ratio was ranked within the range of 3.78±2.09 −13.66±4.05 respectively (Fig.2E, F). C: N ratio contrarily to C: P and N: P showed high value range in the surface and middle depth layers than the last depth (20-30cm). Among the four plant communities within the same depth ranges, the highest C: P and N: P ratio were recorded in *T.quadrispinosa* (81.14±43.88) and *V. natans* (13.7±4.05) respectively and differed significantly (*P* <0.05).

### Leaf, stem and root ecological stoichiometry pattern in four plant communities

The concentration of TOC, TN, and TP among four different plants vary in leaf, root and stem (Fig.3, 4, 5) respectively. Accordingly, the mean TOC, TN and TP concertation in leaf were reported within the range of 92±17.45 g kg^-1^- 197±98 g kg^-1^, 12.88±0.792 g kg^-1^ - 25±2.22 g kg^-1^ and 2.01±0.205g kg^-1^ - 0.6841±0.0263 g kg^-1^ respectively (Fig.3 G, H, I). The highest leaf total carbon (LTC) concentration was measured in *Z. caduciflora* and showed no significance difference (*F*=27.4, *CV*=0.421, *P* >0.05). Among the four plant communities high TN and TP values were measured in *C. schmidtii* leaf and showed significant difference (F=41.96, *CV*=0.26; *P* <0.05; *F*= 33.41, *CV*=0.494, *P* <0.05) respectively. Leaf ecological stoichiometric C: N, C: P, N: P ratios were ranged from 4.92±1.42-9.28±0.827, 82.7±4.606-289±150.59 and 21.4±3.206-96.9±47.57 respectively. Among the four plant communities high C: N and N: P ratios were obtained in *C. schmidtii* and *V. natans* leafs, though, only C: N showed significant difference (*P* <0.05). Highest C: P values were recorded in *Z. caduciflora* leaf and showed significant difference (*F*=48.9, *CV*, 0.728, *P* <0.05). Besides, root total carbon(RTC), root total nitrogen, (RTN) and total phosphorus (RTP) was ranged from 11.16 g kg^-1^ −18.63 g kg^-1^, 1.52g kg^-1^ −5.974 g kg-1 and 0.0615g kg^-1^ −0.644 g kg^-1^ (Fig.4 M, N, O) and showed significant difference (F=53.8, *CV*=0.308, *P*<0.05; F=36.38 *CV*=0.307, *P*<0.05; *F*=59.19, *CV*=0.698, *P*<0.05) respectively. Likewise, R C: P, RC: P, RC: N, and RN: P ratios were obtained within the range of 17.68-109.56, 2.596-9.35 and 4.67-28.95 (Fig.4 P, Q, R) respectively. RC: P, RC: N and RN: P ratios vary significantly (*F*=46.59, *CV*=0.165, *P*<0.05; *F*=62.05, *CV*=0.602, *P*<0.05; *F*=26.52, *CV*=0.103, *P*<0.05) respectively. On the other hand, stem total nitrogen (STN) concertation was measured high in high *C. schmidtii* (Fig.5 T); however, it showed no significant difference (*F*=19.3, *p*>0.05, *CV*=0.58). Among the four studied macrophytes, plant high STN in *C.schmidtii*, whereas high STP was measured in *V.natans* (Fig.5T, U) respectively with slight significant difference (P=0.05).

**Fig. 3.**
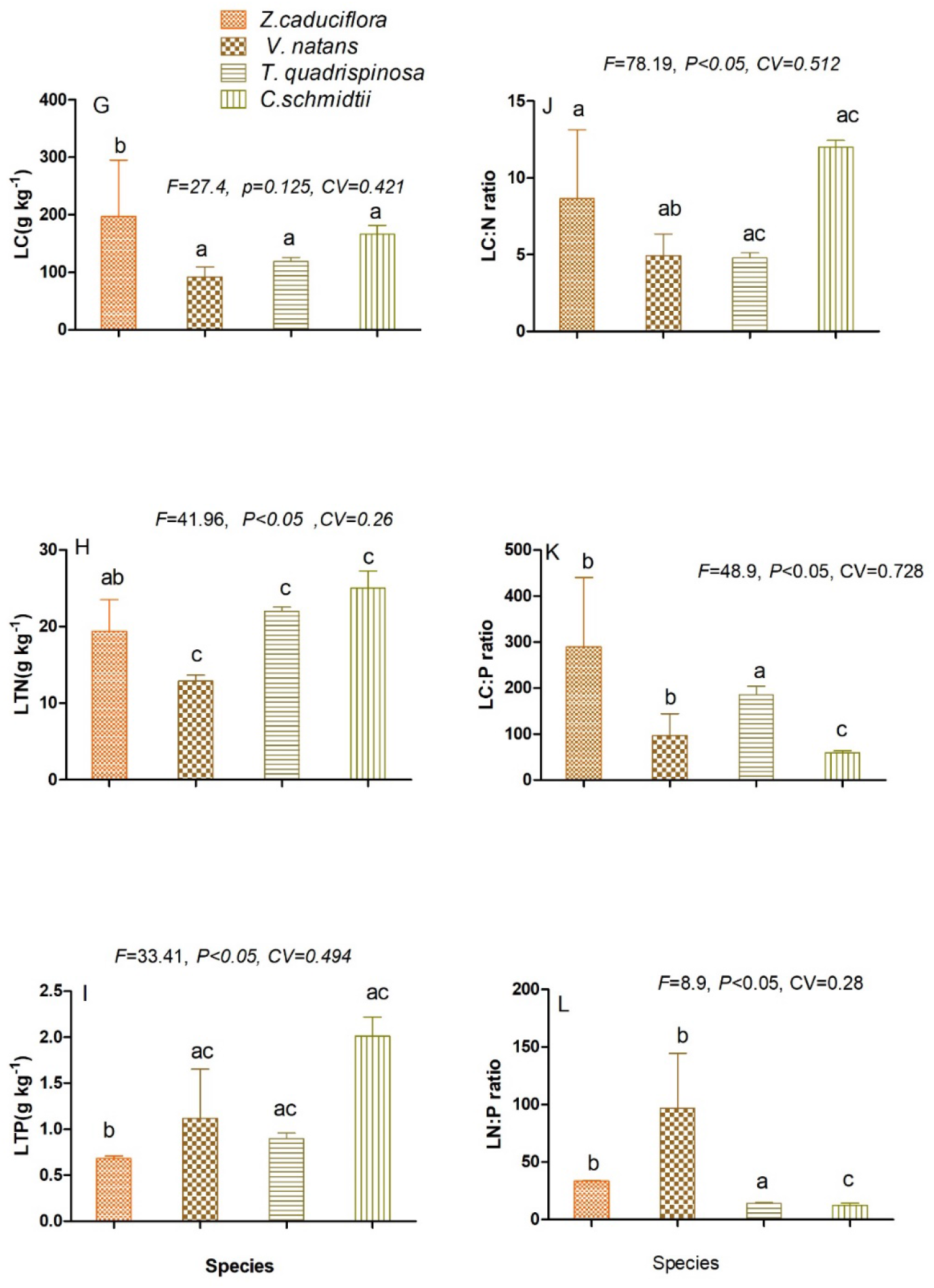
Ecological stoichiometric characteristics of leaf C, N, P and their mass ratio among the four studied macrophytes species. Note: Mean with different letters show significant difference (P<0.05), LSD and Tukey test (2-tailed). LC, leaf total carbon, LTN, leaf total nitrogen, LTP, leaf total phosphorus, LC: N leaf C: N ratio, LC: P leaf C: P ratio, LN: P leaf N: P ratio.

**Fig. 4.**
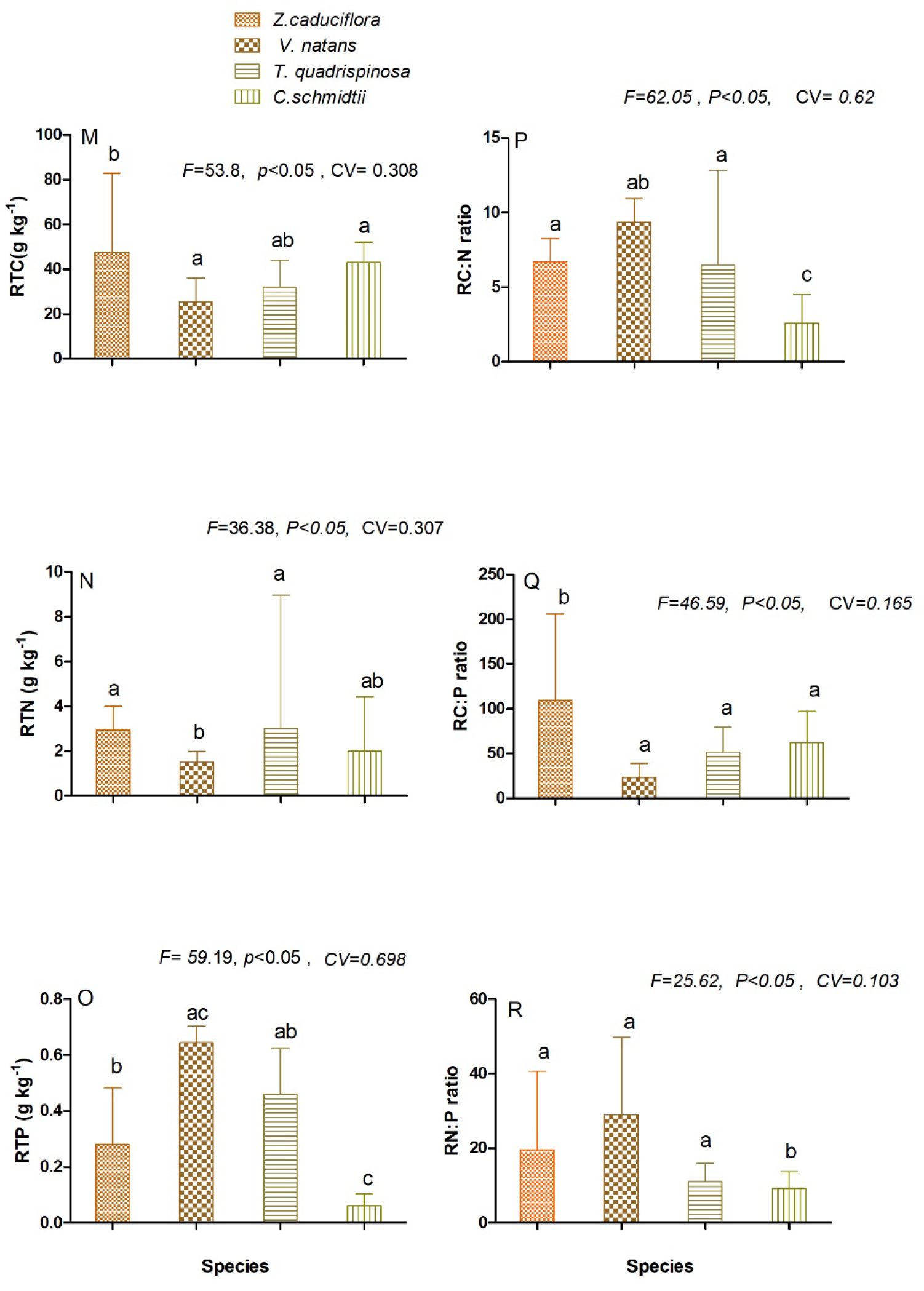
Ecological stoichiometric characteristics of roots C, N, P and their mass ratio among the four studied macrophytes species. Note: Mean with different letters display significant difference (P <0.05), based on LSD and Tukey test (2-tailed). RTC, root total carbon, RTN, root total nitrogen, RTP, root total phosphorus, RC: N, root carbon to nitrogen ratio, RC: P, root carbon to phosphorus ratio, RN: P, root nitrogen to phosphorus ratio.

**Fig. 5.**
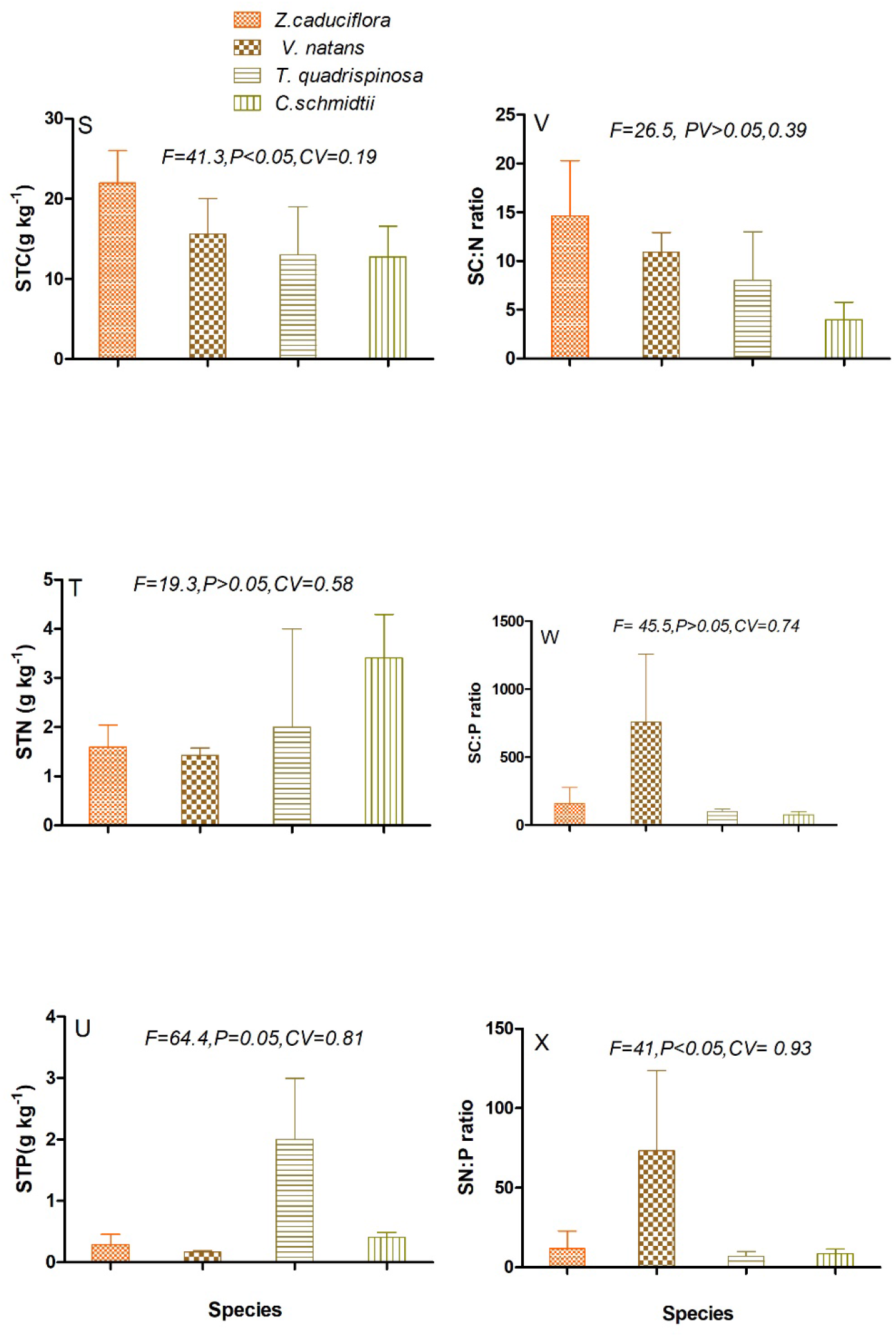
Ecological stoichiometric characteristics of stem C, N, P and their mass ratio among the four studied macrophytes species. Note: Mean with different letters display significant difference (P <0.05), based on LSD and Tukey test (2-tailed). STC, stem total carbon, STN, stem total nitrogen, STP, stem total phosphorus, SC: N, stem carbon to nitrogen ratio, SC: P, stem carbon to phosphorus ratio, SN: P, stem nitrogen to phosphorus ratio.

**Fig.6.**
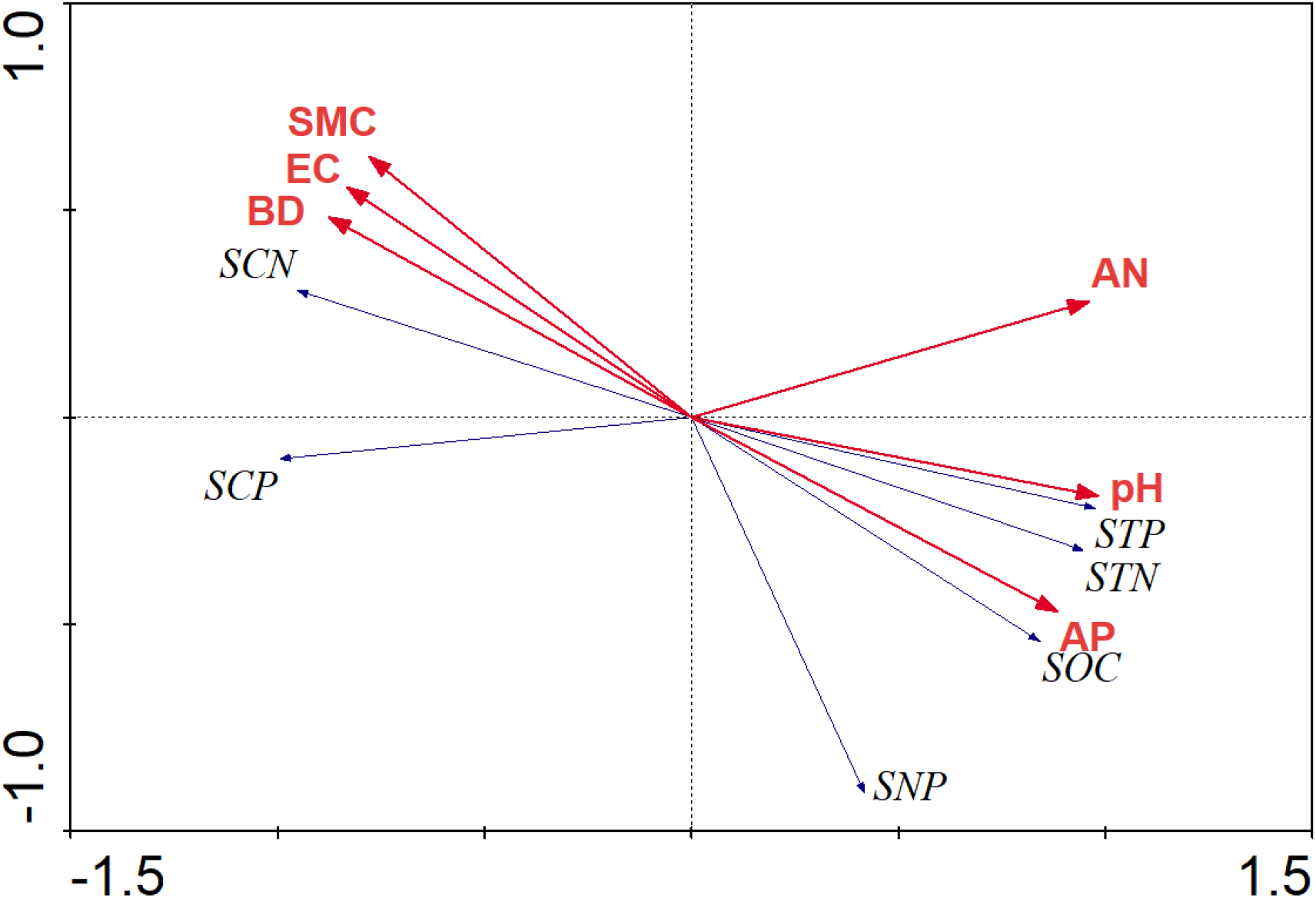
Redundancy Analysis (RDA) result that showed basic soil properties and soil ecological stoichiometric characteristics. Note: In the Box plot SOC indicates, Soil organic carbon, STN, soil total nitrogen, STP, soil total phosphorus, SCN, soil carbon to nitrogen ratio, SNP, soil nitrogen to phosphorus ratio, SCP, soil carbon to phosphorus ratio, EC, electrical conductivity, pH, pH values, SMC, soil moisture contents, BD, bulk density.

### The relationship between soil properties and soil ecological stoichiometry in four plant communities

(Table 1 and 3) showed that there is a considerable significant difference and relationship between soil ecological stoichiometric characteristics and environmental variables respectively. pH values were reported within (5.433±1.10-7.486±0.615) and showed significant difference (*P* <0.05).Furthermore, pH correlated strongly positively with SOC and STN (*P*<0.01), but negatively with STP, SC: N and SN: P ratio (*P*<0.05) (Table 3) respectively. On the other side BD and SMC ranged with (0.904±0.256 −1.48±0.213) and (30.01±14.82-57.38±34.29) respectively (Table 1). Moreover, pH, BD, and SMC were correlated positively with SOC, and STN (*P* <0.01) respectively (Table.3). However, pH negatively correlated with STP, C: N and N: P ratios (*P* <0.05). BD and SMC showed negative relationship with C: N and N: P ratios (*P*<0.05) respectively (Table 3). EC related negative with SOC, STN and STP (*P* <0.05. Available nitrogen (AN) was ranged between (0.351±0.2495-0.602±0.222 mg kg^-L^), whereas available phosphorus was ranged between (24.6±8.63-31.04±7.35 mg kg^-L^) respectively.

**Table 1.** Summary of Redundancy Analysis (RDA) of ecological stoichiometry and soil basic properties

|  | Axis1 | Axis 2 |
| --- | --- | --- |
| Eigenvalues | 0.937 | 0.011 |
| Cumulative percentage variance of species data | 81.6 | 79.2 |
| Species-environment correlations | 0.389 | 0.125 |
| species-environment relation | 80 | 90 |
| *P value* | 0.026* | 0.013* |
| *F ratio* | 3.413 | 0.733 |

### Soil ecological stoichiometric relationship with plant leave, stem and root

As reported in Table 2, SOC content at 0-10cm soil layer negatively correlated with ROC, RTN, RNP, LC, LNP STP, SNP (*P*<0.05), whereas positively related with LTP, LTN(*P*<0.01) and with LC: P (*P* <0.05) respectively. However, SOC at 10-20cm soil layer strongly positively correlated with RTP, LTN, and LC: N ratio (*P*<0.01), whereas negatively with STN (P<0.01) and negatively with STP, SN: P and RN: P ratio (*P*<0.05). Moreover, at 20-30cm it showed strong positive correlation with LTN (P<0.01). On the other side, STN correlated negatively with RTC, LTC, RN: P and LN: P ratios and RTN (*P* <0.05). On the other hand, STN at 0-10cm soil layer strongly positively correlated with LTP, and positively correlated with RN: P and LTC (*P*<0.05). However, negatively correlated with LC: N ratio (*P*<0.05). In the same way STN at 10-20cm soil layer has shown strong positive relationship with LTP, RC: N and RN: P ratio (*P* <0.01) and positively related with RTP (*p*<0.05) but shown negative relation with RC: P and SC: P ratio (*P*<0.05) respectively. At 20-30cm soil layer, STN has also shown strong positive relationship with LTP, STP, RC: N, but negatively with LC: P ratio (*P*<0.01) respectively. With the same manner, STP at different depth layer has also shown positive and negative linear relationship with plant ecological stoichiometric parameters measured in the study. Hence, STP at 0-10cm soil layer related strongly positively with LTN, LTP and RC: N, RC: P RN: P ratio (*P*<0.01) and positively with LN: P and STC (*P*<0.05). Inversely, it showed negative relation with RTC, RTN, STN, STP, SC: P ratio (*P*<0.05). Besides, at 10-20cm soil layer STP was related strongly positively with LTN and RC: P ratio (*P*<0.01) and positively with RC: N and LTP (*P*<0.05). Contrarily, STP with the same depth correlated with LTC and STP negatively (*P*<0.05). On the other side at 20-30cm soil layer it was correlated strongly positively with RC:N and positively with RN:P, LC:P,LN:P ratio and STC (*P*<0.05), however, it indicates strong negative correlation with SC:P ratio (*P*<0.01)and negative correlation with LTC,LC:N, and SC:P ratio (*P*<0.05) respectively.

**Table 2.** Table 2 Basic soil characteristics (Mean±sdav) in different plant communities in Shengjin Lake wetland

| Plant communities | EC (μS/cm) | pH | BD (g·cm<sup>-3</sup>) | SMC (%) | AP(mg kg<sup>-1</sup>) | AN(mg kg<sup>-1</sup>) |
| --- | --- | --- | --- | --- | --- | --- |
| *Z. caduciflora* | 51.05±33.76a | 5.93±0.693c | 1.23±0.219ac | 36.19±9.88a | 0.181±0.1031b | 28.24±9.26b |
| *V. natans* | 64.92±33.96a | 6.72±0.544c | 1.4±0.254a | 57.38±34.29ac | 0.602±0.222a | 31.04±7.35b |
| *T. quadrispinosa* | 48.80±44.7a | 7.486±0.615a | 0.904±0.256ac | 43.81±14.98ab | 0.449±0.2648ac | 24.6±8.63a |
| *C. schmidtii* | 28.29±21.24b | 5.433±1.10c | 1.48±0.213c | 30.01±14.82ac | 0.351±0.2495ac | 27.34±9.75ac |
| Average | 48.02±15.05 | 6.21±0.639 | 1.24±0.252 | 43.19±17.46 | 0.4112±0.1567 | 27.80±2.653 |
| *P* | <0.05 | <0.05 | <0.05 | <0.05 | <0.05 | <0.05 |
| CV | 0.356 | 0.539 | 0.356 | 0.742 | 0.389 | 0.1198 |
| *F* | 28.9 | 61.12 | 73.58 | 56.89 | 29.78 | 89.27 |

Note: Mean with different letter denote significance difference ( $P < 0.05$ ) based on LSD, and Tukey test (2-tailed). EC, electrical conductivity, pH, pH values, BD, bulk density, SMC, soil moisture contents AP, available phosphorus, AN, available nitrogen.

## Discussion

### SOC, TN, TP, and C: N: P distribution patterns among different plants communities

The current study indicates that the stoichiometric characteristics in soil and the plant community has a significant effect on the C: N: P stoichiometry characteristics among the four studied wetland macrophytes plants. Along the vertical gradients, SOC, STN and STP decreased vertically and showed significant difference (*P*<0.05) (Fig.2A, B, C) respectively. This result is in agreement with the previous (Wang J. et al., 2016; Chen et al., 2016) studies. STP found high on the surface than middle and the last depth in this study. Zhang et al., (2005) reported similar result that may indicates P uptake from deep soil to the surface to meet the nutrient requirements may increases on the surface. Soil nutrients have strong effect on plant growth and distribution as well as being the primary source determining the concentration of nutrients in plants (Li LP, et al., 2014; Zhang W, 2015). In this study, the highest nutrient concentration of SOC, STN and STP was observed in *C.schmidtii* and *V.natans* respectively (Fig.2A, B, C) than the rest two species. This may be related with soil moisture contents as the previous study by (Bonetto et al., 1994; Wang et al., 2016) report indicated. Besides, in our study result, soil moisture (SMC) showed positive linear correlation between, SOC, STN and C: P ratio (Table 2), which can confirm this conclusion and coincide with the above two cited study results. Moreover, for high STP measurement on *V.natans* than the rest, community relatively may be related with the returning back of above ground residue to the soil and large number of root decay in wetland ecosystems as indicated by (Chen et al., 2015b). These factors may contribute their own part for high values of TP in this species comparatively. On other hand, high C: P and N: P ratio were found high in *T.quadrispinosa* (81.14±43.88) and *V. natans* (13.7±4.05) respectively and varied significantly (*P* <0.05) (Fig.2E, F). Perhaps this may be in relation with their restoring duration and the plant type or particular species are the main factors in affecting C: P ratio (Mi et al, 2015; Mediavilla et al., 2015). The data presented in this study indicates that, soil stoichiometry at different layers have significantly related with plant stoichiometry (leaf, stem, and root). STN showed significantly positive relationship with leaf total phosphorus (LTP) (*P*<0.001), (Table 2). This result is consistent with (Li et al., 2014) study result that implies soil phosphorus concertation will have direct effect on the photosynthetic active organ to determine not only phosphorus concentration but also nitrogen level too. Moreover, root (C: N: P) also showed positive linear correlation with STP in our study result. Mainly as the previous studies showed this direct relationship might be linked with the genetic and physiological characteristic of the plant that primarily determine elemental concertation and ratio in their tissues (McGroddy et al., 2004; Castle and Neff, 2009). On the other side, stem (C: P: N) ratio negatively correlated with STP at different ranges. This may represents tight coupling coordination between soil nutrient and plant stoichiometric characteristics widely. In addition, the growth rhizomes of the plant themselves and the structure characteristics determine plant tissues nutrients (Baldwin et al., 2006).

### C, N, P and C: N: P variation among plant organs and functional unit in different plant communities

The C, N, P and C: N: P ratio vary among different organs in different plant community are affected by both metabolic demand and the functional differentiation, and organizational structure of plant organs (Minden and Kleyer 2014). Leaf stoichiometry C, N and P paly vital role in analysing composition, structure and functions of a concerned community and ecological systems (Gao et al.,2013). Leaf C found high in Z.caduciflora in this study. This is in agreement with (Baldantoni et al.,2006) that the average of C proportion in emergent plant accounts (45%) and in terrestrial plants (50%) (Agren 2008). TN and TP in different organs was ordered as leaf> root > stem respectively. This may be due to the difference in structure and physiology in different community. Besides, high LTN, however, low N: P ratio was measured in *C. schmidtii* (Fig.3H, L) among the four studied macrophyte plant species. This is supported by (Sardans, and Peñuelas, 2014; Urbina et al. 2017) report that indicate high N and low N: P ratio concentration especially in photosynthetic active organ that species with high growth rates are the best adapted for the environment. This may ascertain the main reason behind the dominance of this marginal plant community especially the beach of the lake, which covers more than 85 % (visual estimation) since 2000 on ward. Thus, a high capacity to retain nutrient in biomass and high nutrient use efficiency can thus be a good traits for plants that grow in wetland areas (Hu et al, 2018) to adapt and increase their size. On the other side, low leaf total carbon (LTC) was measured in *V.natans* (Fig.3G). The possible justification for this result is low leaf carbon is due to less lignin and cellulose content in aquatic plants (Santamaría et al., 2003), because water buoyancy can provide support for aquatic macrophytes, especially for submerged plants. Next to the marginal wetland plant (*C. schmidtii)* which was dominant in our study site, high LTN was measured in *T.quadrispinosa* (Fig.3H) compared to the rest remaining two species. This is consistent with the previous study by (Xie *et al*, 2014) and mainly explained as freely floating macrophytes plant can absorb more N or P from water and sediment through their adventitious root produced from their leaves or stems (Greenway,1997). There is also significant variation among those studied species with regard to P and N concentration in both stem and root parts and as well C:N:P ratios. For instance, high P and N: P ratio was measured in *V.natans* root whereas least in *C. schmidtii* (Fig.4O, R) respectively. In general, this may shows that, submersed macrophytes plants are able to uptake nutrients both from water column by their leaves or stems and from sediments by their roots or rhizoids (Madsen and Cedergreen, 2002; Cao et al., 2011) these can contribute these two nutrients become high relatively in this species. In ecological studies, leaf N: P ratio was considered as an important index to identify limiting nutrient elements (Güsewell, 2004, Wu et al., 2012a). Particularly for wetland ecosystems when N: P <14 plant growth is limited by N, while when N: P >16 plant growth limited by P, and when N: P 14<N: P>16 the growth limited by both N and P (Wu et al., 2012a). In the present study, leaf N: P ratio found > 16. This revealed that P nutrient element was found as a limiting nutrient element than N in our study.

### The relationship between soil ecological stoichiometry and environmental variables

Basic biogenic elements in soil ecosystems such as C, N and P were closely related with soil physicochemical properties (Feng *et al*., 2017, Zhang *et al*, 2017). In this same way, in study, soil pH values ranged between (5.433±1.10 −7.486±0.615), (Table 1) and positively correlated with SOC and STN (*P*<0.01), (Table 3). This may indicate more or less the range of wetland pH values scale that range around 6.5 −7.5 with few exceptions in general (Gambrel, 1994). In addition, this is in agreement with the previous results reported by (Lu *et al*., 2018). However, pH values showed inverse relationship with STP, C: N and C: P ratios (*P*<0.05), (Table 3). This is inconsistence with (Feng *et al*., 2017; Lu et al., 2018) study report which contradict the present study. This may indicate that high pH values in the soil can restrain total nitrogen decomposition (Sun *et al*., 2017). EC ranged (28.29±21.24-64.92±33.96) (Table 1) and correlated negatively with SOC, STN, and STP (*P*<0.05) respectively (Table 3). This is coincide with (Yu *et al*., 2016), but inconsistent with (Tashi *et al*., 2016) pervious study report. SMC correlated positively with SOC, STN (*P* <0.01) and with STP and C: P ratio (*P*<0.05) (Tabel.3) respectively. This is in agreement with (Yang *et al*., 2016; Zhang *et al*., 2017; Lu *et al*., 2018) studies. Most previous studies indicated basic properties for instance pH, SMC, BD can regulate the dynamic change of C: N and C: P ratios (Sun *et al*., 2017). In addition, SMC positively correlated with C: P (*P* <0.05).This may indicates high SMC can increases organic phosphorus and raise C: P ratio in wetland ecosystems Zhang (*et al*., 2017). However, negatively correlated with C: N ratio. This coincides with (Mayor *et al*., 2017; Lu *et al*., 2018), however, it contradicts (Sun *et al*., 2017) pervious study result. BD correlated with SOC and STN strongly positively (*P* <0.01) and positively with STN, STP, and C: N ratio (*P* <0.05) respectively. This result is inconsistent with (Lu *et al*., 2018).

**Table 3.** Pearson correlation analysis result between soil ecological stoichiometry and plant nutrients in roots, stems and leaves along the depth gradients

|  | Soil C |  |  | Soil N |  |  | Soil P |  |  |
| --- | --- | --- | --- | --- | --- | --- | --- | --- | --- |
|  | 0-10 cm | 10-20 cm | 20-30 cm | 0-10 cm | 10-20 cm | 20-30 cm | 0-10 cm | 10-20 cm | 20-30 cm |
| Root C | -0.982* | -0.773* | 0.226 | 0.181 | 0.999* | 0.876* | -0.834* | 0.346 | -0.252 |
| Root N | -0.998* | 0.343 | -0.999* | -0.952 | 0.885 | 0.238 | -0.998* | 0.772 | -0.431 |
| Root P | 0.788 | 0.096 | 0.710** | 0.984* | 0.738* | -0.171 | -0.268 | 0.566 | 0.694 |
| Root C: N | 0.691 | 0.926 | 0.625 | 0.603 | 0.851** | 0.912** | 0.895** | 0.277 | 0.998** |
| Root C: P | 0.114 | 0.133 | 0.061 | -0.893* | -0.819** | 0.805 | 0.944** | 0.819** | -0.263 |
| Root N: P | -0.999* | -0.565* | -0.733* | 0.965* | 0.571** | 0.784* | 0.964** | 0.999* | 0.423* |
| Leaf C | -0.883* | -0.856* | 0.080 | 0.805* | 0.738* | 0.632* | -0.392 | -0.783* | -0.850* |
| Leaf N | 0.922** | 0.457 | 0.962** | 0.146 | 0.939 | -0.119 | 0.890** | 0.257** | 0.487 |
| Leaf P | 0.827** | 0.999** | 0.828** | 0.815** | 0.849** | 0.784** | 0.237 | 0.673* | -0.366 |
| Leaf C: N | 0.793 | 0.944** | 0.492 | -0.893* | 0.177 | 0.219 | 0.368 | -0.391 | -0.282* |
| Leaf C: P | 0.794* | 0.159 | 0.224 | 0.115 | 0.191 | -0.962** | 0.253 | 0.368 | 0.106* |
| Leaf N: P | -0.936* | -0.757* | 0.138 | 0.083 | -0.053* | 0.259 | 0.673* | 0.287 | 0.132* |
| Stem C | 0.36 | 0.07 | 0.921 | 0.083 | -0.421* | 0.290 | 0.726* | 0.285 | 0.233* |
| Stem N | 0.168 | -0.80** | -0.331* | 0.249 | 0.17 | 0.223 | -0.377* | 0.044 | -0.648* |
| Stem P | -0.599* | -0.446* | 0.305 | 0.264 | 0.191 | 0.771** | -0.703* | -0.331* | 0.054 |
| Stem C:N | 0.825 | 0.408 | 0.296 | 0.128* | 0.47 | 0.026 | 0.101 | 0.210 | 0.149 |
| Stem C:P | -0.882 | 0.799 | 0.666 | 0.456 | 0.327 | 0.703* | -0.622* | 0.218 | -0.982** |
| Stem N:P | -0.582* | -0.804* | 0.739 | 0.249 | 0.553 | 0.225 | 0.015 | 0.327 | 0.064* |

Note: Correlation with \* star shows significant difference ( $P < 0.05$ ), (2-tailed analysis).
Correlation with \*\* star shows significant difference ( $P < 0.01$ ) (2-tailed analysis).

**Table 4.**
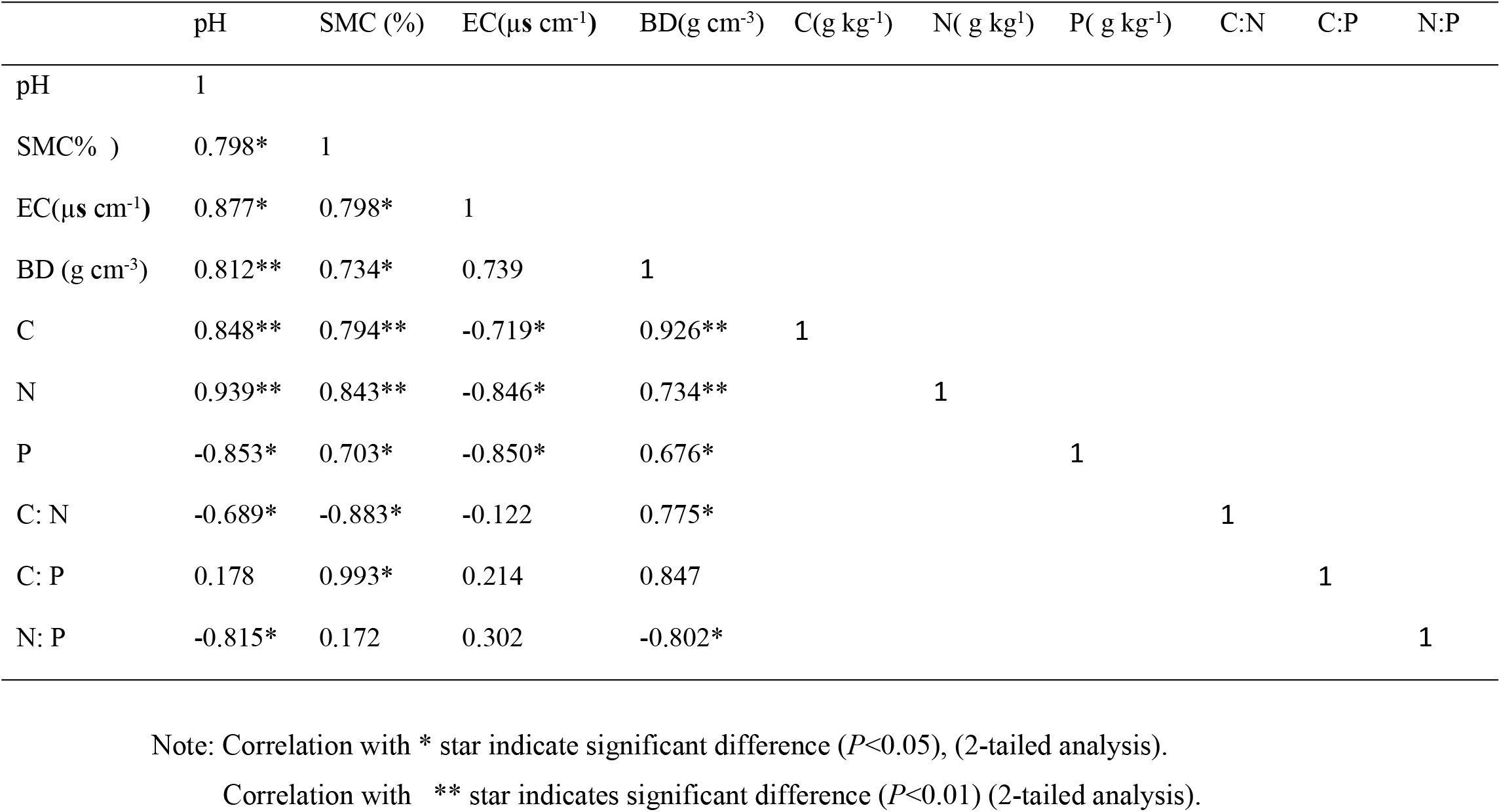
Pearson correlation analysis result between soil ecological stoichiometric characteristics and environmental variables

| | pH | SMC (%) | EC($\mu\text{s cm}^{-1}$) | BD($\text{g cm}^{-3}$) | C($\text{g kg}^{-1}$) | N( $\text{g kg}^{-1}$) | P( $\text{g kg}^{-1}$) | C:N | C:P | N:P |
| --- | --- | --- | --- | --- | --- | --- | --- | --- | --- | --- |
| pH | 1 |  |  |  |  |  |  |  |  |  |
| SMC% ) | 0.798* | 1 |  |  |  |  |  |  |  |  |
| EC($\mu\text{s cm}^{-1}$) | 0.877* | 0.798* | 1 | | | | | | | |
| BD ($\text{g cm}^{-3}$) | 0.812** | 0.734* | 0.739 | 1 | | | | | | |
| C | 0.848** | 0.794** | -0.719* | 0.926** | 1 |  |  |  |  |  |
| N | 0.939** | 0.843** | -0.846* | 0.734** |  | 1 |  |  |  |  |
| P | -0.853* | 0.703* | -0.850* | 0.676* |  |  | 1 |  |  |  |
| C: N | -0.689* | -0.883* | -0.122 | 0.775* |  |  |  | 1 |  |  |
| C: P | 0.178 | 0.993* | 0.214 | 0.847 |  |  |  |  | 1 |  |
| N: P | -0.815* | 0.172 | 0.302 | -0.802* |  |  |  |  |  | 1 |

Note: Correlation with \* star indicate significant difference ( $P<0.05$ ), (2-tailed analysis).
Correlation with \*\* star indicates significant difference ( $P<0.01$ ) (2-tailed analysis).

### Conclusion and implications

For health and sustainability of ecosystems vegetation restoration can play significant roles in distribution and accumulation of soil and plant stoichiometric characteristics. The C, N, P and C: N: P ratio vary among different organs in four plant community has been investigated after the lake wetland vegetation began restoring overtime. The resulted showed variation among those studied plant communities. These resulted from the functional differentiation, life forms and organizational structure of plant organs. High LTN and low N: P ratio was measured in *C. schmidtii*. This is related with active organ that provides best to adapt for the environment. This may ascertain the main reason behind the dominance of this marginal plant community especially at the beach of the lake. High P and N: P ratio was measured in *V.natans* root due to their ability to uptake nutrients both from water column by their leaves or stems and from sediments by their roots or rhizoids. Soil pH and available phosphorus (AP) were found the potential variables that affect soil ecological stoichiometry from the RDA analysis result.

## Acknowledgement

This research was funded by the National Natural Science Foundation of China (NFSC) [31800346]. The authors would like to thank all the reviewers for their valuable and constructive comments in improving the original manuscript.

